# Make it grow, *Pseudozyma aphidis* extract promotes plant growth

**DOI:** 10.1101/2025.01.12.632596

**Authors:** Anton Fennec, Neta Rotem, Maggie Levy

## Abstract

Increasing global population demands higher agricultural yield while avoiding the use of harmful products in agriculture. The use of beneficial micro-organisms is an Eco-friendly approach to reduce the use of fertilizers and plant growth promoting (PGP) products. This study is focused on the fungus *Pseudozyma aphidis (P. aphidis)*, a yeast like dimorphic fungus beneficial to plants and related to the phytopathogenic fungus *Ustilago maydis* that causes corn smut disease. Our lab isolated a unique *P. aphidis* strain (PA12) that has shown significant plant protection and plant growth activity. We extracted and screened the active PGP fraction from PA12, calibrated treatment dosage and application method. Tomato, corn and melon plants treated with PA12 extract exhibited; increased germination, shoot length, biomass and higher yield. Our results demonstrate the potential of *P. aphidis* extract application as an eco-friendly PGP agent that can contribute to reduction of harmful agricultural products usage.

## Introduction

The rapid growth of global population and the constant demand for higher agricultural yield requires intense use of fertilizers and pesticides. A study by Australia’s National Science Agency revealed rapid buildup of Nitrous Oxide (N2O) in the atmosphere and ice cores in the past 2000 years (CSIRO, 2022). Fertilizers are a major source of N2O which is considered a greenhouse gas that contributes to the increasing climate changes effect (Rochette et al., 2008). Between 200 A.D to 1800 A.D N2O was at a relatively stable concentration, with the increasing population, industry and agriculture by 2022 it increased to 335ppb, a 24% increase in only 222 years of rapid anthropogenic activity (CSIRO, 2022). Another example is the environmental effect of extensive fertilization that increases the rate of emerging dead zones. Between the years 1960 to 2008, over 400 new dead zone systems were reported in comparison to ∼50-56 that were known before that time (Diaz & Rosenberg, 2008). Thus, reduction of intense fertilizers and pesticides use have become more than just a necessity but rather a new challenge.

There are various approaches to deal with this challenge without severe compromise on crops and yield. The use of beneficial micro-organisms (BMO’s) is an Eco-friendly approach that promotes reduced fertilizers and potentially harmful plant growth promoting (PGP) products application (Yadav et al., 2020). Various fungal and bacterial species are studied for their beneficial effects. Micro-organisms employ several known methods to enhance plant growth. The most common and known PGP modes of action include the following: phosphate solubilization, nutrient uptake, plant protection, stress tolerance, phytohormones and nitrogen fixation (Singh, 2018). Some fungi and bacteria can produce the phytohormone Indole acetic acid (IAA), similar to that of plants. Both plant-beneficial and pathogenic micro-organisms are known to possess this capability (Chung et al., 2003, Sun et al., 2014). Additionally, glycolipid surfactants produced by several microorganisms demonstrated PGP activity and theorized as bioactive compounds with a direct mode of action (Marchut-Mikołajczyk et al., 2021a).

Although the application of some BMO’s is already used in agricultural practice there are many complications that follow. Deferent species require diverse conditions thus the application of live BMO’s is somewhat limited by geography and climate (Ruocco et al., 2011). Additional problem with this practice is the uncertainty of how the BMO established on the host plants and how persistent its beneficial activity might be (Blum et al., 2011). The presence of live BMOs may also push towards faster arms race evolution of the pathogenic microorganisms (Simberloff & Stiling, 1996).

The most common approach to establish a long-lasting BMO growth on the plants is by repeated application of new BMO inoculum (Blum et al., 2011). Our approach takes us in a completely different direction by eliminating the need to maintain the BMO in the field and creating more repeatable and uniform effect by application of eco-friendly BMO based compounds. The effect of BMO based extracts of algae and several microorganisms were tested mostly on juvenile plants and tomatoes (Metwally et al., 2022). Thus, much of the information that is available is partial and limited to juvenile plants and often model organism plants. Most studies on PGP related fungi are conducted on arbuscular mycorrhizal fungi and some are done on epiphytic and endophytic fungi but to lesser extent (Bacon & White, 2016; Elsherbiny et al., 2023). Because majority of these studies are limited to juvenile plants there is scarce information on the fungal PGP effects on mature plants biomass and size (Nimsi et al., 2023). Furthermore, very little is known on their effects on the yield and crop quality. Often treatments that enhance plant growth and crops may reduce the crop quality parameters such as total soluble solids (TSS), an indicator for sugar, titratable acid (TA) for sourness indicator. Additional parameters that are often affected by treatments enhancing the crop yield include titratable nitrogen (N) that acts as indicator for nutritional value and produce firmness (Iyer et al., 2012). The produce taste is another major parameter of produce quality that often suffers a drawback from treatments that enhance plant growth and crop yield (Tandon et al., 2003). In fact, it has become one of the major complaints among consumers that the taste of agricultural produce is bland, and the cause seems to be our prerogative to grow more food and faster (Estabrook, 2012).

*P. aphidis* is a yeast like dimorphic fungus related to the phytopathogenic fungus *Ustilago*. Unlike its relative, *P. aphidis* is not only non-pathogenic but rather beneficial to the host plant (Barda et al., 2015; Buxdorf et al., 2013; Gafni et al., 2015). We isolated a unique *P. aphidis* strain (PA12) that has shown significant anti-fungal and anti-bacterial activity in previous studies in our lab (Srivastava et al., 2021). PA12 exhibits antagonistic, parasitic activity against powdery mildew (Gafni et al., 2015). *P. aphidis* secreted lytic enzymes that activate programmed cell death (PCD) when paired with in live *B. cinerea* (Buxdorf et al., 2013; Calderón et al., 2019). Tomato plants treated with live PA12 cell culture exhibited enhanced plant growth and improved yield (Shoam et al., 2022). Cucumber seedlings were also examined and has shown enhanced growth after being treated with live PA12 cultures, but did not reach the fruiting stages (Shoam et al., 2022). Early work on PA12 extracts conducted in our lab has demonstrated bioactive and plant protective qualities of PA12 secretions but the PGP aspect was not examined nor characterized (Harris & Levy, unpublished data). Several Pseudozyma species have shown PGP activity, mostly in seedlings and juvenile plants. *P. antarctica* produces and secretes itaconic acid (methylidene succinic acid) that increases plant nutrient uptake (Levinson et al., 2006; Zhao et al., 2022). *P. aphidis* isolates were found to produce ammonia (NH_3_) and auxin. Most the PGP capable isolates were fund capable of calcium phosphate tribasic solubilizing activity; thus, they can enhance nutrient uptake to the host plant (Fu et al., 2016). For the most part, studies on the Pseudozyma PGP and biocontrol effects of live cultures and extracts were conducted on several cultivars, their focus remained on the biocontrol aspects(Golubev et al., 2001, Metwally et al., 2022). A study by Metwally et al, (2022) on fungal extracts from Trichoderma cultures have shown that C-phycocyanin from the extract enhanced tomato germination and seedlings growth (Metwally et al., 2022). The majority of the work focused on plant protection, hence the necessity to extend our study on the PGP activity aspect of *P. aphidis*. We extract the bioactive PGP fraction secreted by the PA12 and applied it on the seeds and plants as they grow. Thus, we eliminated the dependency of establishing a live BMO colony on the plants and get more consistent treatment with known and repeatable concentration with the PGP extract.

Our PA12 extract acts as an eco-friendly PGP agent that in addition to increasing plant growth, biomass and induce early flowering, it also enhanced the produce yield and quality. Our data circumrotates the characterization of the PA12 extract activity on the full process of growing crop baring plants relevant in agriculture from germination to harvest and characterizing the effect on the crop quality. These results demonstrate the potential of compounds extracted from *P. aphidis* to contribute to green agriculture and global food security and encourage a further study of the mechanisms involved in its activity.

## Materials and Methods

### PA cultures and plant material

Corn, Raymon melon and M-82 tomato seeds were sterilized in 3% bleach solution for 5 min, then washed 3 times with sterile distilled water, and placed at 4 °c overnight before germination. The treated seeds were coated by dipping them in 3mg/ml extract and then dried, the control seeds were coated with water. The seeds were planted in vermiculite and kept in a growth chamber at 22°c with 10/14 light dark cycle. The seedlings were grown in the growth chamber for 3-7 days, then transferred to a larger pot (30cm) with commercial potting soil medium and moved to a clear walled green house (natural day-night cycle). Conditions in the greenhouse were 24°c, 60% RH and automated irrigation (100% flow, 10 min-morning and evening). After the growth of the first two leaves, the treated plants were sprayed with 3mg/ml extract on by bi-weekly basis until the fruit initiation growth stage, the control plants were sprayed with water.

### PA extract preparation, screening and dosage

PA12 isolate used for extract preparation was grown on Potato Dextrose Agar (PDA) Petri plates at a 28°c incubator for 10 days. 2 PDA plates were collected and transferred to a 250ml round culture flask with 50ml Murashige and Skoog (MS) liquid medium. The flasks were grown in a dark, orbital shaker incubator at 28°c, 175 rpm for 10 days. The cultures from the flasks were taken for extract preparation. The extracts were made directly from the full culture media with the cell suspension. PA flasks were extracted with 4 equivalents of organic solvents gradient from hydrophilic (Ethanol) to hydrophobic (Hexane) on an orbital shaker at 300rpm for 30-180min. The organic phase was collected, and solvents were evaporated with a Buchi Rotary evaporator system (R-215). Dry extract was collected, weighed and resuspended in a 40eq aqueous solution (70/30%, DW/extraction solution) **(Table S1)**.

The solvent media combination extracts were screened for optimal PGP performance using a bioassay. Col-o arabidopsis seeds were treated with the combinations extract, germinated and grown for 10 days. Root and shoot length were measured in order to identify the optimal culture medium and solvent combination **(Table S1)**. Next, we examined the top PGP preforming extract (Hexane/Acetone) for antimicrobial activity by using antibiotic paper disc assay. *Botrytis cinerea* and *Agrobacterium tumefaciens* cultures were grown for 24 hours on PDA and LB plates, then a Whatman paper disc containing 1-50 mg/ml extract was placed on the centre of the culture plate. The inhibition halo was examined for 5 consecutive days **(Table S2)**. This allowed us to screen for an extract that has only PGP activity.

The extracted fractions were screened for PGP activity by coating Col-0 arabidopsis seeds with crude extract fraction and growing them on MS solid medium plates. The treated 7-day old seedlings root and shoot length were evaluated for the selection of the optimal dosage and solvent and culture media combination **(Table S1)**.

Pharmacological assays were made on 15-30 seeds per treatment (0-10 mg/ml extract) the dried extract re-suspended in ethanol, up to volume of 400 µl. Lethal Dose (LD), Effective Dose (ED) and Toxic dose (TD) are the doses that have lethal, positive and toxic effect on portion of the population. These parameters were calculated based on the common pharmacological principals established by Miller and Tainer (1944) and revised in 2009 (Miller & Tainter, 2009). We used a population of treated and control Col-0 arabidopsis seeds and seedling in LD and LD50, that dose that kills 50% of calculation. The number of treated germinating seeds and seedling morality for up to 10 days after germination was normalized according to the Log method (Singh et al., 2020). ED and ED50, the effective dose that has a positive effect on 50% of the population was calculated by incorporating germination, root and shoot length of the germinated seedlings. The data was standardized with Log method and the ED50 principals (Barreto et al., 2002). TD is the toxic dose and TD50 is the dose that is toxic/harmful but not lethal to 50% of the population. It was calculated as the inverted values of ED (Jones, 1990).

The best PGP extract solvents combination was hexane 70%: acetone 30%, diluted to a 3 mg/ml concentration for application according to pharmacological test and it did not exhibit any antimicrobial activity in vitro.

### Screening for application method

Following the determination of extract conditions and dosage, we examined parameters by creating a multivariate correlation matrix of PGP activity and selected those with value of r^2^>0.8 in correlation to other PGP parameters. The Chosen parameters were used for testing extract application methods of seed coating, spraying the extract on germinated plants every two weeks, and combined seed coating and spraying **(Fig. S2)**. This screening allowed us to further tune the extract and its application method to an optimum performance.

### Germination, flowering, plant and leaves size, number of leaves and biomass measurement

Germination was evaluated by percentage of the germinating seedlings per total sowed seeds and evaluated after 10-14 days after no new germinating seedlings sprouted. Flowering was evaluated from the time stamp a plant started to develop flowers until reaching full mature bloom and evaluated as percentage of flowering plants out of all the plants. Shoot height was measured on 5-week-old plants and mature plants at harvest time. leaves length and the number of leaves of 5-week-old plants was manually evaluated. Mature plants size was evaluated by Fiji ImageJ. Dry weight of mature plants and 100 corn kernels was evaluated by fully drying in a 45°c oven.

### Tomato ripening and firmness

Tomato fruit ripening was determined visually based on standard color ripening scale based on scale used by Takizawa et al (Takizawa et al., 2014). For the evaluation of fruit ripening, 5 stages of fruit maturation were used; green-unripe green fruit, breaker-tomatoes that started to develop slight red color towards ripening, turning-the stage that a large portion of the fruit starting to turn to pink/red color, light red-whole fruit is pale red, red-fully ripe red fruit. The ripening rank was scored visually and all the harvested tomato fruits were examined per plant per treatment. The fruit firmness of the harvested tomatoes was determined with a Shore analog durometer (Fei et al., 2018). The firmness scale of the fruit was categorized according to a range of durometer units (DU); rank 1-very soft 0-10 DU, rank 2-soft 11-20 DU, rank 3-flexible 21-30 DU, rank 4-firm 31-50 DU and rank 5-hard 51-100 DU.

### TSS, TA, Nitrogen content and taste test

Tomato juice total soluble content, was acquired by the method of Yadav et al. (Yadav et al., 2021). Ripe fruit juice, 3ml of 5 fruit was filtered with 5 layers of cheese cloth, diluted by 1:1 ratio with DW. 1ml was taken and measured in °Brix with a fruit juice refractometer. Total titratable acid and titratable N content of the tomato juice was measured with the method used by the teams of Heeb and Majidi (Heeb et al., 2005; Majidi et al., 2011). For determination of TA, 5ml of tomato juice was titrated with 0.1N NaOH to pH 8. Then, the titrated samples were treated with formaldehyde and further titrated with 1 N NaOH to pH 8 for total N content. The TA results expressed in citric acid equivalent (eq) mg/ml. The taste test of mature ripe tomato fruit was based on the method used by (Heeb et al., 2005). Fruit from mature treated and control plants were harvested, washed and labelled with random letters. In the tasting tests, 32 people were presented with a questionnaire ranking sweetness, sourness, texture, color and overall preference. The parameters were ranked from 1-lowest, less preferable to 10-highest, most preferable.

### Statistical analysis

Statistical analysis was conducted with SAS JMP 11software for; multiple comparisons, multivariate, correlation, matrix analyses and students t-test were used according to the experiment at α=0.05.

## Results

### PA extract calibration pharmacological assays and characterization

This study aimed to extract and isolate the PA secreted bioactive PGP fraction in order to characterize its activity by application on crop plants. We screened crude extract fractions in a large range from hydrophobic to hydrophilic affinity from cultures in MS and PDB media and found that the optimal extract conditions are MS medium cultures extracted with Hexane:Acetone 70/30% v/v **(Table S1)**. Then we tested our fraction for antimicrobial activity and found that it did not have any direct activity against *B. cinerea* and *A. tumefaciens* **(Table S2)**. Thus, we successfully separated the compounds responsible for PGP from those with antibiosis activity. Pharmacological tests of the PGP extract on Col-o seeds and seedlings show that LD50 of our extract is 6.3 mg/ml, TD50 is 4.5 mg/ml and ED50 is 3.5 mg/ml. We decided to use the extract at concentration of 3 mg/ml, that allows ∼70% ED with optimal repeatable results (**Fig. 1**).

**Figure 1.**
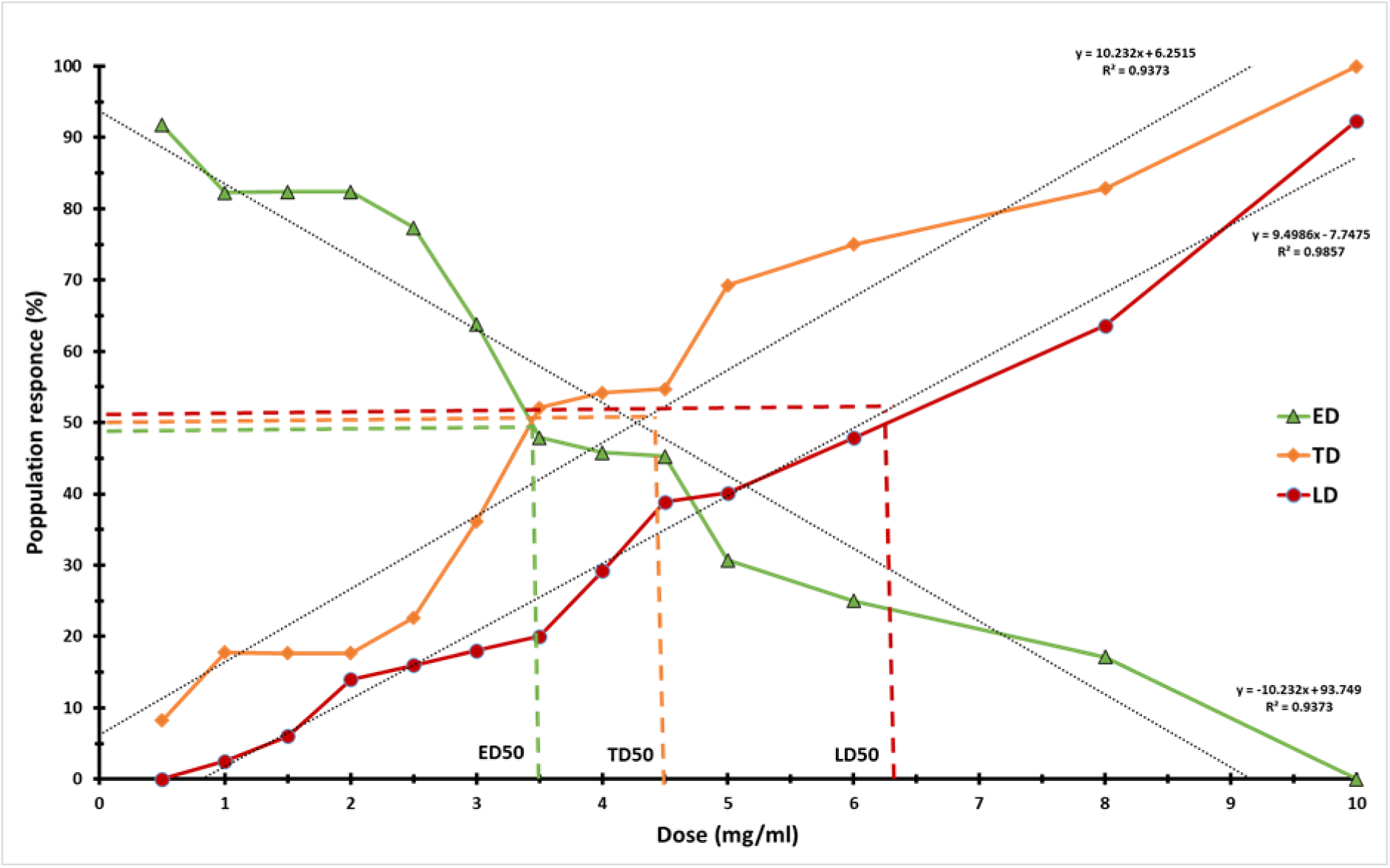
Pharmacology and optimal extract dose. The effective dose (ED, green), toxic dose (TD, orange) and lethal dose (LD, red). LD is calculated from the population portion of germinating seeds and seedlings survivability (up to 10 days post germination). ED is calculated from the portion of seeds population that germinated, their survivability and positive effect on root and shoot length (standardized with the Log method). TD was calculated as an inverse trend of ED (see methods). LD50 is the dose that is lethal to 50% of the population, TD50 is a dose with toxic effect on 50% of the population and ED50 is the dose with positive effect on 50% of the population. Dashed lines represent: LD50=6.3mg/ml, TD50=4.5mg/ml, ED50=3.5mg/ml.

### Calibration of application method

Looking at a single parameter alone can be highly inaccurate when screening for an optimal application method with multiple effects. Multivariate correlation matrix between PGP parameters of all the extract application methods (seed coating, spray and combined) and control, assists in selecting the parameters and their combination that contribute the most to PGP activity. The parameters that had high correlation (r^2^ value) indicate parameters that can be used as determinants for growth promoting activity for further selection of optimal application method **(Fig.S1 a & b)**. Thus, application with significantly higher performance in parameters that correlate for growth promoting activity were selected for further work. We concluded that seed coating combined with bi weekly spray and seed coating alone had the best performance and combined seed coating and spray had a slight edge in regards to crop ripening (Fig. S1c).

### PA extract effect on juvenile plants germination and growth parameters

In order to characterize the effect of PA12 extract on crop plants (tomato, corn and melon) during their juvenile stage, we examined the germination and additional growth parameters of treated 5-week-old plants compared to the untreated control (**Table 1**). All plants treated with PA12 extract had significantly higher germination rate compared to the control (**Table 1**). Tomato plants have shown 18% increase in germination rate following PA12 extract treatment. Lesser, but significant effect was observed on treated corn and melon seeds germination with an increase of ±7% (**Table 1**). We also found that the coated seeds of all the plants germinated 2 to 3 days earlier than the corresponding control seeds (data not shown). Treated juvenile plants to 5-week-old plants grew significantly longer shoot and leaves (**Table 1**). Tomato shoot was 9.5% longer, melon was 12.2% longer and corn was 21.4% longer than control (**Table 1**). Treated melon plants had significantly 24.3% more leaves compared to control. No effect on number of leaves was observed on corn and tomato plants (**Table 1**). Melon leaves length was 8% longer, corn leaves 13.8% and tomato leaves were 34.6% longer in treated plants compared to control.

**Table 1.** Physiological parameters of different crop plants after treatment with PA12 extract. Mean and standard error accompanied by connecting letters represent significant deference calculated with students t test, α=0.05; n=100 (germination), 20 (shoot length, leaves length and number of leaves).

| Plant | Group | Germination (%) | Shoot length (cm) | leaves length (cm) | Number of leaves |
| --- | --- | --- | --- | --- | --- |
| Corn | Control | 92.16 ± 1.09 b | 15.86 ± 0.43 b | 47.8 ± 2.48 b | 6 ± 0 |
|  | Treatment | 100 ± 0 a | 20.2 ± 2.15 a | 55.46 ± 0.89 a | 6 ± 0 |
| Melon | Control | 86.7 ± 2.23 b | 22.46 ± 1.26 b | 8.54 ± 0.34 b | 3.1 ± 0.16 b |
|  | Treatment | 93 ± 1.09 a | 25.6 ± 1.48 a | 9.29 ± 0.37 a | 4.1 ± 0.16 a |
| Tomato | Control | 77.7 ± 7.8 b | 28.3 ± 0.7 b | 11.7 ± 2.4 b | 9 ± 0.7 |
|  | Treatment | 96 ± 2.6 a | 31.3 ± 0.5 a | 17.9 ± 2.9 a | 9.2 ± 0.9 |

### PA extract effect on flowering time

Flowering development and time to full bloom of mature plants was evaluated (**Fig.2**). Treated plants reached full flowering 1 to 2 weeks earlier than their control counterparts (**Fig.2**). Treated melon and tomato plants started flowering and achieved full bloom 1 week before the control plants. The treated melon plants started flowering 2 weeks before the control plants but reached full bloom only one week sooner compared to the control (**Fig. 2**).

**Figure 2.**
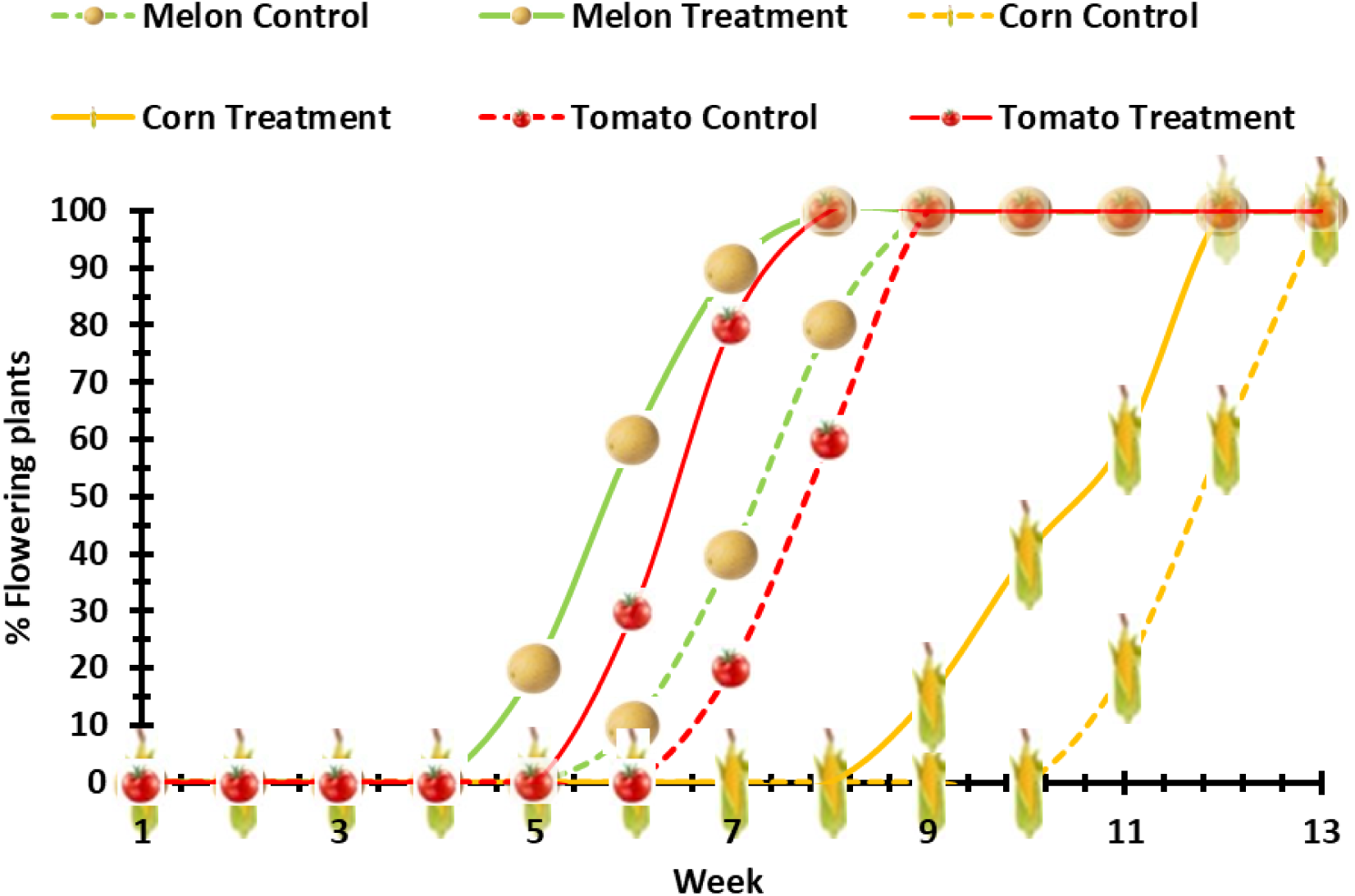
Flowering time from start to full bloom. Time stamps of plants of all cultivars (melon-green, tomato-red and corn-yellow), continuous lines represent treated plants and dash lines represent the control plants. Data is shown in weeks from flower formation on the plants to full blossom of all plants for each cultivar. The results are shown in % of the flowering plants of each cultivar vs all the plants per given cultivar.

### PA extract effect on mature plant growth and yield

Upon plants maturation, we examined the effect of PA12 extract treatment on plant growth, mass and yield of the three crop plants; tomato, melon and corn. Shoots of mature plants treated with PA12 extracts were significantly longer in all the crop plants compared to the untreated control. Shoot height of tomato was 22% higher, melon was 36% higher and corn was 11.4% higher compared to the control (**Fig.3a, b& c**, and **Fig.3d, e& f**). Additionally, the fresh and dry weight of the treated plants was also significantly higher than that of the control plants (**Fig.3g, h& i**). Fresh weight of treated tomato plants was higher by 29.4% and melon was 48.5% higher compared to the control (**Fig.3g & i**). No significant effect on corn plants fresh weight was observed (**Fig.3h**). The dry weight of treated plants of all the crops has substantially increased; tomato by 42.7%, corn by 30.1% and melon by 64.2% as compare to control (**Fig.3j, k& l**).

**Figure 3.**
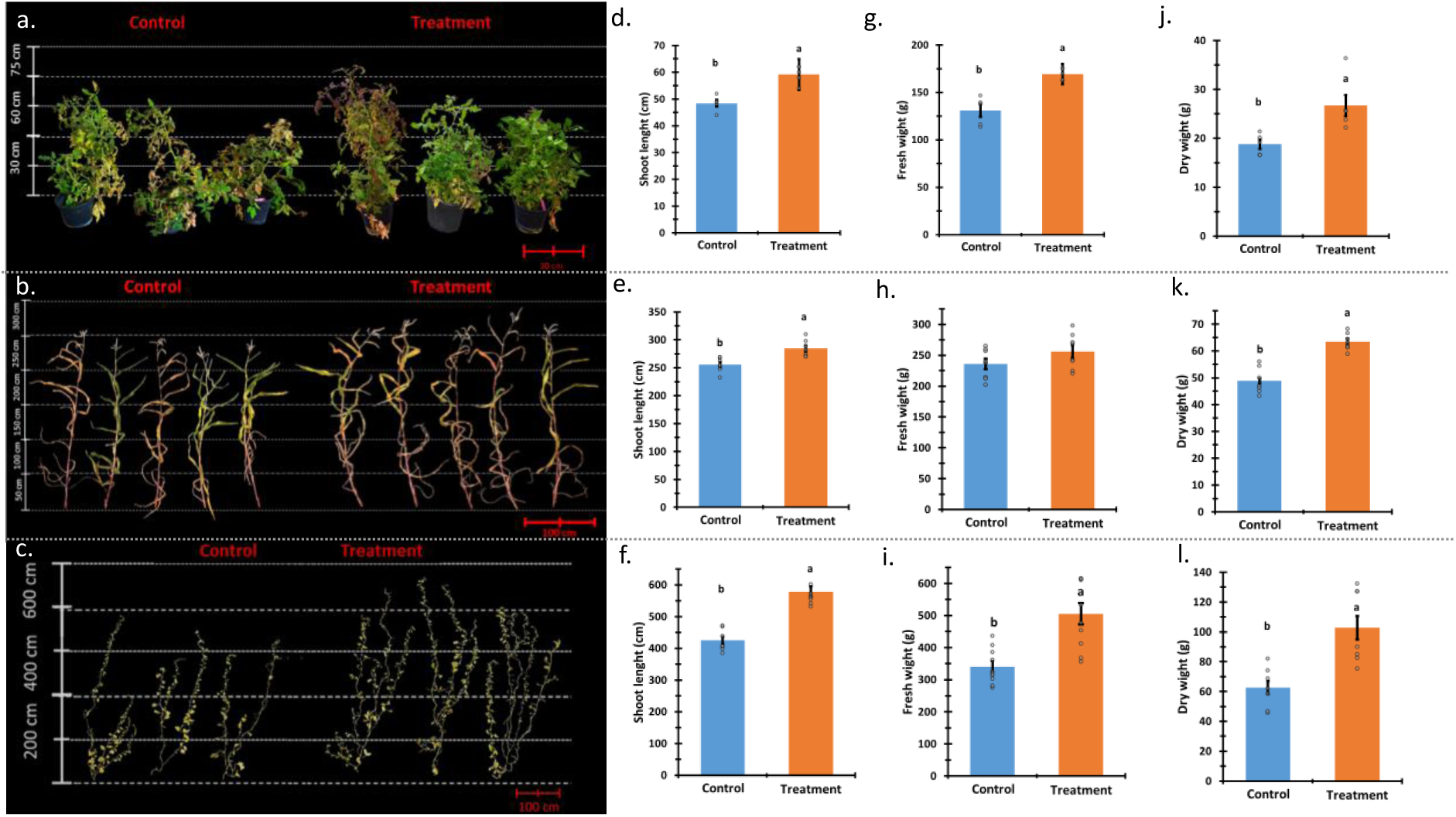
Plant height and mass. Representative tomato, corn and melon plants pictures with scale measure (a to c accordingly). Shoot length of the plants of all the cultivars; d-tomato, e-corn and f-melon. Fresh and dry weight of the plant stalks; tomato-g & j, corn-h & k, melon-i & l (accordingly). Blue bars represent the average data of control group and the orange bars represent treated plant group. Mean and standard error accompanied by connecting letters represent significant deference calculated with students t test, α=0.05; for shoot length n= 5 (tomato plants), 7 (corn and melon plants). For fresh and dry weight n= 5 (tomato), 8 (corn) and 10 (melon plants).

Once the plants matured, we characterized the PA12 extract treatment effect on yield. The harvested crop was examined for yield and ripening parameters (**Fig.4**). Treated plants produced distinguishably higher yield compared control. Tomato plants yielded fruits in total weight of 61.4% higher and 18.4% more tomato fruit per plant compared to the control. Additionally, the fruit from the treated tomato plants ripened sooner (300% more ripe fruit), and the ripening was more uniform compared to the control (**Fig.4a, d, g &J**). Treated melon plants produced average fruit weighing 429.4% more than that of control and larger in diameter by 148%. An average treated melon plant produced 90.3% more fruit par plant compared to the control (**Fig.4b, e, h &k**). Average fresh and dry weight of 100 kernel corn from treated plants was 15.9% and 19% higher than the control (accordingly). The treated corn plants also produced 88.6% more cobs per plant (**Fig.4c, f, i &l**).

**Figure 4.**
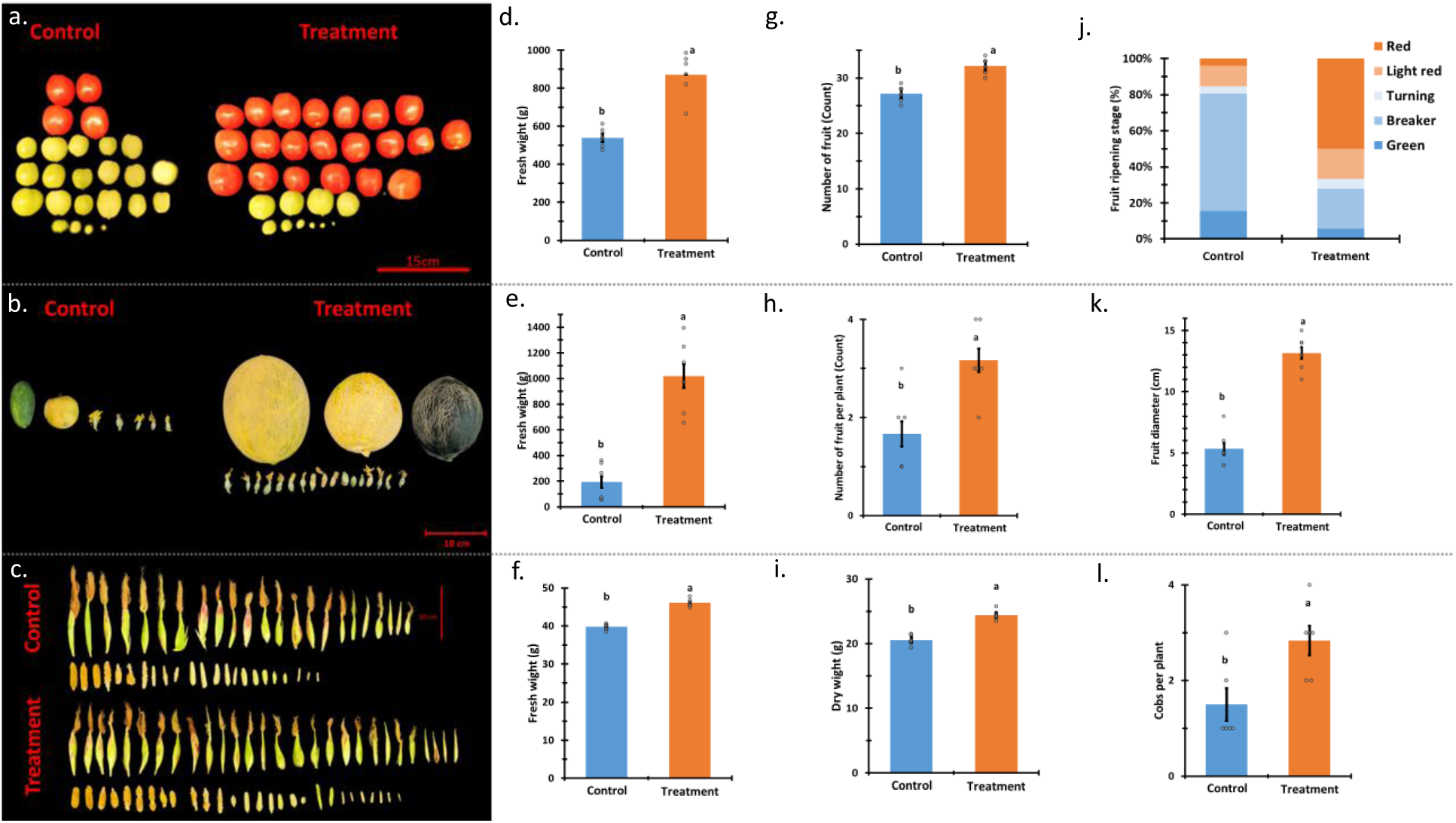
Crop, yield and ripening. Representative tomato fruit (per plant), melon (per plant) and corn cobs (collected from 10 plants) pictures with scale measure (a to c accordingly). Fresh average weight of tomatoes per plant-d, melon fruit-e and 100 corn kernels-f. Number of fruit from tomato-g, melon plants-h and 100 corn kernel dry weight-i. Tomato ripening fruit by stage represented in percentage from total number of fruit (ranking system in methods section)-j, average melon fruit size expressed in diameter-k and number of cobs per corn plant-l. Blue bars represent the average data of control group and the orange bars represent treated plant group. Mean and standard error accompanied by connecting letters represent significant deference calculated with students t test, α=0.05; for all the parameters n= 6.

Next, we examined the effect of the PA12 extract treatment on tomato fruit quality and fruit taste (**Fig.5**). Fruit of treated plants had significantly higher TSS by 30.6%, 72.7% higher TA and 345% more titratable nitrogen content compared to the control (**Fig.5a, b & c**). Additionally, to the nutritional parameters, fruit firmness of from threated plants was also significantly higher by 35.7% (**Fig.5d**). Next, a taste test was conducted in order to examine the treatment effect on the flavor of the treated tomatoes (**Fig.5e**). Fruit of treated plants received significantly higher testers score in all parameters (sweetness, acidity, aroma, color and texture). The color and texture score were higher by 2 ranks and sweetness, acidity and aroma higher by 1 score rank. The overall choice score of tomatoes from treated plants was 2 ranks (20%) higher compared to the untreated control (**Fig.5e**).

**Figure 5.**
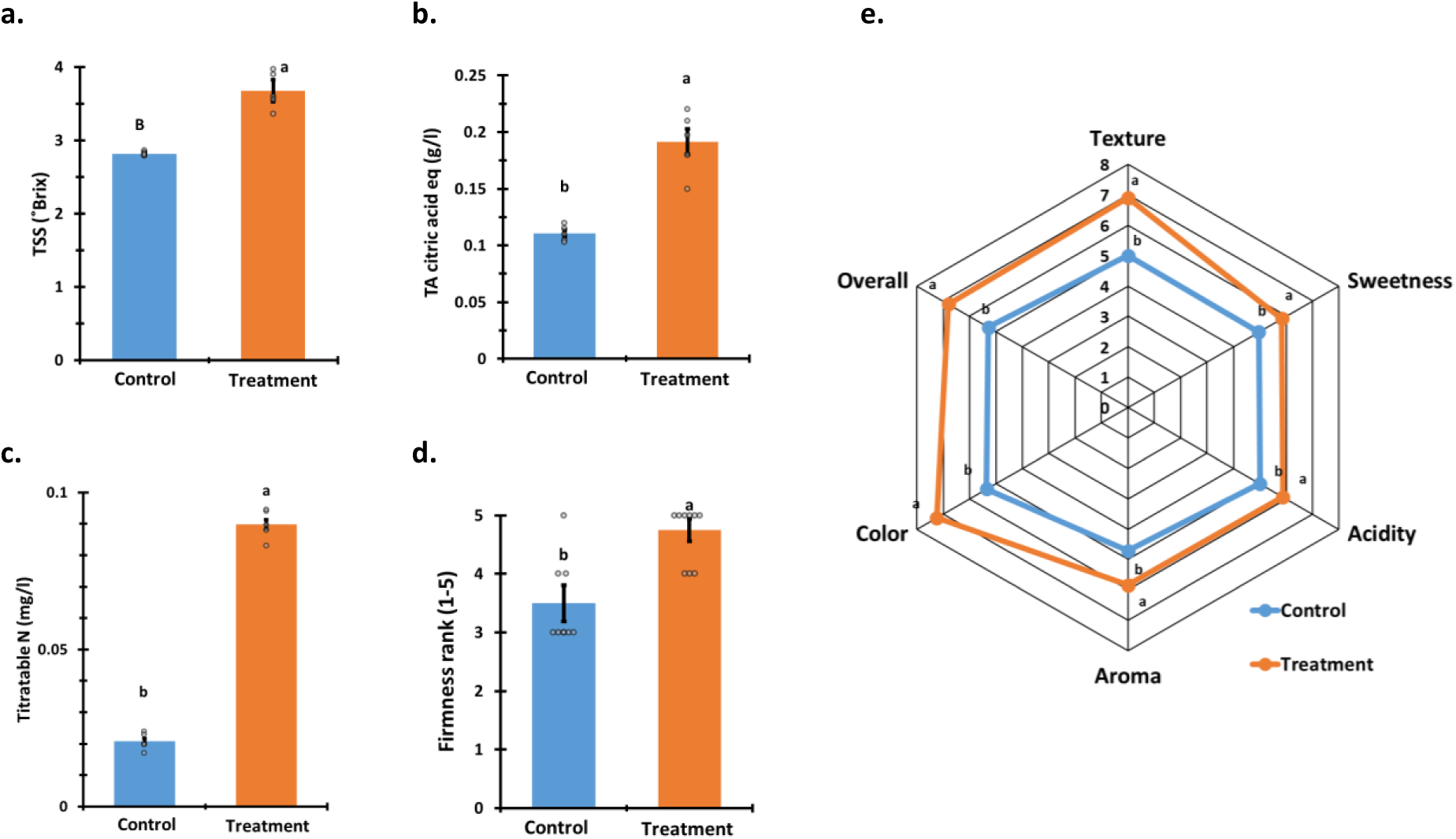
Tomato fruit quality. Fruit quality was expressed in TSS-a, TA-b, titratable nitrogen content (N)-c, fruit firmness index (as described in methods section)-d and a taste test (described in methods)-e. blue bars represent the average data of control group and the orange bars represent treated plant group. Mean and standard error accompanied by connecting letters represent significant deference calculated with students t test, α=0.05; for TSS, TA and N n= 5. For Firmness and taste test n= 9 & 32, accordingly.

## Discussion

Although *P. aphidis* is known to have PGP activity, the application, effects on yield and crop quality are not well characterized. In previous work in our lab, treatments with live PA12 cultures enhanced plant growth and increased the yield of M-82 Tomato plants (Shoam et al., 2022). Similar observations regarding PGP activity of Pseudozyma species by various research groups are reported in the literature (Boekhout, 2011; Fu et al., 2016; Levinson et al., 2006). In this study we aimed to extract and isolate the PA secreted bioactive PGP fraction in order to characterize its activity by application on crop plants.

We found that the optimal extract conditions to investigate the PGP activity were an extract from MS media culture, extracted with Hexane:Acetone **(Table S1)**. Our screening method coupled with PGP activity bioassay have shown that the extract we selected did not have any antibiosis properties in vitro **(Table S2)**. Thus, we successfully separated the compounds responsible for PGP and active antibiosis activity. This approach allowed us to reduce the interference from antibiotic activity and characterize the PGP activity of PA12 extracted secreted compounds. Bioactive molecules are known to have a positive effect in low concentration and lethal in high concentration (Puga et al., 2012). We determined the optimal beneficial concentration of the PA12 extract in range between 3 to 4.5 mg/ml (**Fig. 1**) and found that the optimal application method is seed coating combined with bi-weekly spraying (**Fig. S1**). Those application methods are standard application methods in agricultural practice (Verkleij, 1992).

In our experiments, extract concentration of 3.5 mg/ml was used for seeds coating and bi-weekly spraying followed by measuring different physiological effects on several crop plants. After treatments with PA12 PGP fraction we found tremendous effects on plant growth and yield of all three crop plants we examined: tomato, corn and melon. Flowering time of treated plants of all the crops was one to two weeks sooner than that of the control plants in both flowering start and in time of full bloom. Tomato and melon plants flowering came one week early and corn started flowering two weeks ahead of its control (**Fig.2**). All the treated plants reached full bloom one week before the control plants. To our best knowledge this is the first report of early flowering induced by *P. aphidis* or its extract. Majority of the work on PGP is done on arbuscular mycorrhizal fungi (AMF), and the data that supports early flowering induction by fungi comes from AMF (Sohn et al., 2003). Early flowering is a possible precursor to early fruit set and development that can shorten growing season and might have significant implementation for fast goring crops (Nocker & Gardiner, 2014). Our results have shown that the PA12 extract treatment indeed worked as a precursor for early fruit set and development, in particular in melon plants.

The shoot length of all the crop plants treated with PA12 extract was significantly higher compared to the control plants. Juvenile treated plants of all crops were higher by 10-33%. Shoots of mature melon treated plants were most effected by the treatment and were higher by 30% than control plants. Mature tomato and corn shoot length was also significantly affected by the PA12 extract treatment but to a lesser extent of 10 to 20 % longer (**Fig.3**).

The fresh weight of tomato and melon plants was significantly higher by 30% and 50% than that of the control accordingly. While fresh weight of corn did not show any significant alteration as compared to control (**Fig.3**). The dry plants weight of treated tomato and melon was significantly almost 2 times higher than that of the control, the corn dry weight was 50% higher than the control (**Fig.3**). The deference in corn fresh and dry weight in regards to the control can be attributed to higher water content in the control plants. Lower water content accompanied by higher biomass was shown to be a trait allowing grain producing plants to maintain water potential and photosynthesis in drought resistant conditions (Internat. Rice Res. Inst., 1982). This might indicate a possibility of the PA12 extract treatment as drought protectant. The increased plant biomass is also in an increasing demand for fuel and animal feed uses (Spiertz & Ewert, 2009).

Our result demonstrates how PA12 extract treatment increased the plant biomass, that can also contribute to its increasing demand as animal feed.

An increase in plant biomass, size and early flowering by plant growth stimulants alone does not necessarily indicate positive effect on the crop and the yield, as demonstrated by Zhang and Whiting (2011) (Zhang & Whiting, 2011). Our results have shown that in addition to enhanced plant growth, our PA12 extract had a positive effect on crop yield of all the treated plants. Fruit yield and quality of tomatoes from treated plants were unexpectedly higher. The fruit from PA12 treated plants ripened sooner (over 60% ripe fruit) and the plants produced nearly 20%?? more fruit (**Fig.4**). We observed similar trend in previous studies using live PA12 cultures for treatments on microtome tomatoes (Shoam et al., 2022). While the fruit on the control melon plants barley started to grow, the treated plants already had large mature fruit (**Fig.4**). Melon plants treated with PA12 extract produced fruit that weighed five times more than that of the control plants, each plant produced twice more fruit and its size (diameter) was more than twice than that from the control plants. Corn kernels of treated plants weighed 17% and 20 % more than those of the control (fresh and dry accordingly) and each corn plant produced nearly twice more cobs (**Fig.3**). Increasing the produce of grains has an enormous significance in establishing food security. The possibility to achieve it with an eco-friendly BMO based approach can reduce the agricultural foot print on the environment due to the large area required to grow grain crops (Peng et al., 2023).

Increase in crop and yield is often accompanied by relinquished produce quality including negative effect on flavor (Todeschini et al., 2018). But it was also shown that grape berries fruit treated with PGP micro-organisms on ripening fruit can improve their quality (Ding et al., 2019). Surprisingly, we found that the fruit of plants treated with our PA12 extract contained more natural sugars and higher quantity of nitrogen-based compounds. Thus, this fruit are sweeter and with higher nutritional value (**Fig**.**5**).

Fruit of the treated tomato plants were 30% firmer as compared to the control (**Fig**.**5**). The connection between firmness, fruit quality and shelf life are established in many cultivars (Matas et al., 2009). Hence, the PA12 extract treatment possibly also enhances the produce shelf life and may aid in food loss reduction.

Although fruit quality as measured by sugar content, soluble acids, nutrient values and fruit firmness has good correlation to flavor sensory, it does not always represent the flavor quality of produce (Iyer et al., 2012; Tandon et al., 2003). To complement the fruit quality data, our taste tests have shown that the flavor of the tomato fruit of treated plants was 10 to 20% time higher ranked in all taste parameters (**Fig**.**5**). Flavor parameters of tomatoes from treated plants (sweetness and acidity) and aroma were ranks slightly higher compared to the control while the texture, color and overall preference was scored two ranks higher (**Fig.5**). These results demonstrate the ability of PA12 extract to increase the produce, its quality and even to improve the flavor.

In this study, we have calibrated a method and conditions to extract the bioactive PGP fraction from PA12 cultures. We identified the optimal dosage and calibrated the application method, characterized its effect on plant growth and demonstrated the effect on crop yield, quality and flavor. Our method of extract preparation should be further tuned to purify and identify the PGP compounds and the mechanisms involved in its activity. This study elaborates the PGP activity of PA12 extract application on crop plants, and demonstrates the ability of compounds secreted by PA12 to enhance plant growth, increase the yield and the produce quality while also improving the flavor. Thus, this study can help to promote BMO’s Eco-friendly agricultural application and aid in global food security.

## Acknowledgements

Deepest gratitude to Inbal Sharaby for assistance in the corn and melon experiments, and for assistance in maintaining the PA12 cultures. Thanks to Gilli Breur and Yahav Bracha for their help in the tomato experiments and PA12 cultures maintenance.

## Statements and Declarations

### Conflict of Interest

There are no competing interests/conflict of interests, financial and non-financial regarding any aspect of this paper with any of the authors involved in the making of this paper directly and indirectly.

### Funding

Funding for this project was provided by the Israeli Ministry of Agriculture, grant number-12-02-0043.

### Author contribution

AF-experimental design and execution and data analysis, NR-technical and experimental advice and support, ML-experimental design, advice and project layout. The manuscript was written by AF and ML. All authors read and approved the manuscript.

## Notes

### Competing Interest Statement

The authors have declared no competing interest.

