## supplementary figures and methods for "Make it grow, *Pseudozyma aphidis* extract promotes plant growth"

| Treatment | Root length (cm) | Shoot length (cm) |
|---|---|---|
| Cont DW | 1.21 ± 0.03 h | 0.34 ± 0.1 h |
| Cont PDB | 1.21 ± 0.02 h | 0.35 ± 0.01 h |
| Cont MS | 1.18 ± 0.03 h | 0.41 ± 0.02 fg |
| PDB Ethanol | 1.5 ± 0.07 cd | 0.55 ± 0.03 abc |
| MS Ethanol | 1.31 ± 0.05 g | 0.38 ± 0.02 gh |
| PDB Chlorophorm | 1.5 ± 0.04 cd | 0.45 ± 0.02 ef |
| MS Chlorophorm | 1.44 ± 0.04 f | 0.53 ± 0.02 cd |
| PDB Et Ace | 1.48 ± 0.02 cde | 0.6 ± 0.01 a |
| MS Et Ace | 1.45 ± 0.03 ef | 0.5 ± 0.1 de |
| PDB Acetone | 1.45 ± 0.03 f | 0.54 ± 0.02 bcd |
| MS Acetone | 1.48 ± 0.02 def | 0.52 ± 0.02 cd |
| PDB Hex Acet | 1.52 ± 0.02 c | 0.58 ± 0.02 ab |
| MS HEX Acet | 1.56 ± 0.02 a | 0.6 ± 0.01 a |
| PDB Hexane | 1.3 ± 0.01 g | 0.5 ± 0.01 de |
| MS Hexane | 1.61 ± 0.01 a | 0.56 ± 0.02 abc |

**Table S1**. Root length and shoot length of Col-0 arabidopsis plants treated with 3-4 mg/ml PA extracts of deferent solvent and culture media combination (MS and PDB). The solvents were selected by their hydrophilic/phobic characteristics from ethanol to hexane. Root and shoot length measurements were obtained from 7-day old treated seedlings. Mean and standard error accompanied by connecting letters represent significant deference calculated with students t test, $\alpha=0.05$; n=50 plants per media/solvent combination treatment.

| Extract | Concentration | Bo.5 halo | A. tumefaciens halo |
|---|---|---|---|
| MS HEX Acet | 1mg/ml | 0 ± 0 | 0 ± 0 |
| | 5mg/ml | 0 ± 0 | 0 ± 0 |
| | 10mg/ml | 0 ± 0 | 0 ± 0 |
| | 25mg/ml | 0 ± 0 | 0 ± 0 |
| | 50mg/ml | 0 ± 0 | 0 ± 0 |

**Table S2**. PA extract test for antimicrobial activity. MS hexane acetone solvent combination tested for antimicrobial activity against *B. cinerea* and *A. tumefaciens* invitro. Agrobacterium and Botrytis were grown on solid medium for 24 hours and then a disc with extract was placed in the centre of the petri dish. The disk was loaded with 1mg/ml to 50 mg/ml extract and the inhibition halo was measured after 5 days.

a

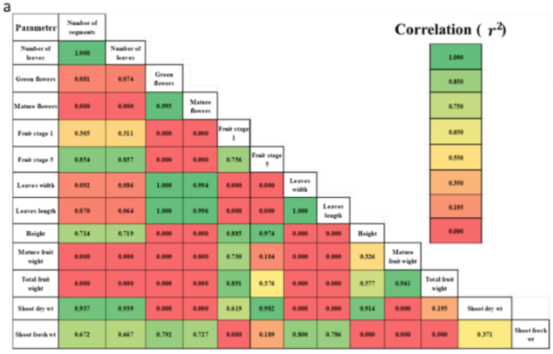


b

| | | | | | |
|---|---|---|---|---|---|
| Number of leaves | Number of segments | 1.000 | Total fruit wight | Mature fruit wight | 0.961 |
| Leaves width | Green flowers | 1.000 | Shoot dry wt | Number of leaves | 0.939 |
| Leaves length | Green flowers | 1.000 | Shoot dry wt | Number of segments | 0.937 |
| Leaves length | Leaves width | 1.000 | Shoot dry wt | Height | 0.914 |
| Leaves length | Mature flowers | 0.996 | Total fruit wight | Fruit stage 1 | 0.891 |
| Mature flowers | Green flowers | 0.995 | Height | Fruit stage 1 | 0.885 |
| Leaves width | Mature flowers | 0.994 | Fruit stage 5 | Number of leaves | 0.857 |
| Shoot dry wt | Fruit stage 5 | 0.982 | Fruit stage 5 | Number of segments | 0.854 |
| Height | Fruit stage 5 | 0.974 | Shoot fresh wt | Leaves width | 0.800 |

C

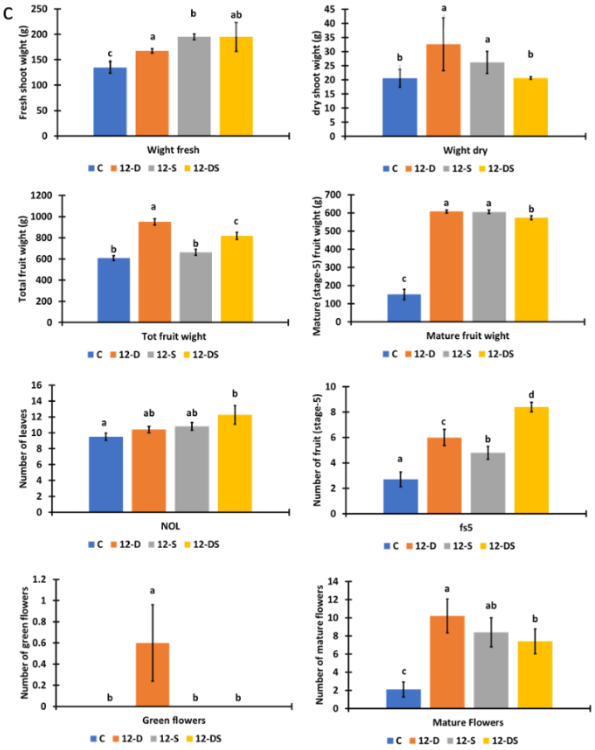


**Fig**. **S1** Multivariate correlation matrix of all the PA12 treatments. a. Multivariate matrix, green indicates high correlation between parameters across all treatments and red indicates low to no correlation between the parameters across all treatments. b. table of parameters with high correlation across all treatments that indicate growth promoting activity. Parameters with R^2 values higher than 0.8 were selected. c. Comparison of treatment methods on all significant parameters, the column bars represent the average values, error bars represent Standard Error. Connecting letters are Significance as obtained by Students T-test, α=0.05.
